# Establishment of an Inhalation Administration Non-invasive Murine Model for Rapidly Testing Drug Activity against *Mycobacterium tuberculosis*

**DOI:** 10.1101/2024.02.27.582260

**Authors:** Xirong Tian, Yamin Gao, Wanli Ma, Jingran Zhang, Yanan Ju, Jie Ding, Sanshan Zeng, H.M. Adnan Hameed, Nanshan Zhong, Gregory M. Cook, Jinxing Hu, Tianyu Zhang

## Abstract

The efficacy of many compounds against *Mycobacterium tuberculosis* is often limited when administered via conventional oral or injection routes due to suboptimal pharmacokinetic characteristics. Inhalation delivery methods have been investigated to achieve high local therapeutic doses in the lungs. However, previous models, typically employing wild-type *M. tuberculosis* strains, were intricate, time-consuming, labor-intensive, and with poor repeatability. In this study, we developed an autoluminescence-based inhalation administration model to evaluate drug activity by quantifying relative light units (RLUs) emitted from live mice infected with autoluminescent *M. tuberculosis*. This novel approach has several improvements: it eliminates the need for anesthesia in mice during administration and simplifies the instrument manipulation; it is cost-effective by utilizing mice instead of larger animals; it shortens time from several months to 16 or 17 days for obtaining result; it is non-invasive by measuring the live RLUs of mice; up to six mice can be administrated daily and simultaneously, even 2-3 times/day; results are relatively objective and repeatable minimizing human factors. Proof-of-concept experiments demonstrated that inhalable rifampicin, isoniazid, and ethambutol showed anti-*M. tuberculosis* activity at concentrations as low as 0.5, 0.5, and 0.625 mg/mL, respectively, as evidenced by comparing the live RLUs of mice. Furthermore, consistency between RLUs and colony-forming units of the lungs reaffirms the reliability of RLUs as an indicator of drug efficacy, highlighting the potential of this approach for accurately assessing anti-*M. tuberculosis* activity *in vivo*. This autoluminescence-based and non-invasive inhalation model offers a substantial reduction in the time, effort, and cost required for evaluating the efficacy of screening new drugs and repurposing old drugs *in vivo* via inhalation administration.

## INTRODUCTION

*Mycobacterium tuberculosis*, primarily transmitted through respiratory droplets, is the causative agent of human tuberculosis (TB) and is responsible for the highest mortality rate attributed to a single infectious agent before the 2019 coronavirus pandemic (McQuaid et al., 2021, Shah et al., 2022). The emergence of drug-resistant *M. tuberculosis* strains presents substantial public health challenges and economic burdens (Dousa et al., 2020, Hameed et al., 2018, McGrath et al., 2014, Shoen et al., 2018). Therefore, there is an urgent need for innovative *in vivo* drugs screening and evaluation tools to identify efficacious antimicrobial agents for TB control. Conventional *in vivo* efficacy evaluation experiments for anti-TB drugs often employ oral or various injection routes of administration (Namvarpour et al., 2019, Yu et al., 2020, Yu et al., 2021). However, these methods fail to specifically target the lungs and may induce significant toxicity in other organs during absorption, transport, and metabolism (Xu et al., 2019, Yoon et al., 2017, Zhao et al., 2023). Over several decades, inhalation therapy, delivering high local therapeutic doses to the lungs, has emerged as a preferred treatment method for respiratory diseases (Barnes et al., 2015, Garcia-Contreras et al., 2007, Gonzalez-Juarrero et al., 2012, Hickey et al., 2016, Sanchis et al., 2013). However, previous attempts to develop inhalable anti-TB drug administration models have encountered several challenges, including the time-consuming process of counting colony-forming units (CFUs), the complexities associated with anesthesia during administration, the difficulty in instrument operation, and the poor repeatability of experimental results (Garcia-Contreras et al., 2010, Verma et al., 2013).

Meanwhile, most existing drugs evaluation models mentioned above are based on wild-type *M. tuberculosis* and require several months for obtaining results, as it takes 3-6 weeks for the *M. tuberculosis* to form visible colonies on agar plates. Recently, some researchers have introduced autoluminescent *M. tuberculosis* into the process of drugs or vaccine evaluation (Yang et al., 2015, Zhang et al., 2012). The *luxAB* genes encode the enzymes that catalyze the reaction resulting in the emission of blue-green light, and *luxCDE* genes are responsible for the synthesis and recycling of the aldehyde substrate (Hakkila et al., 2002). Light production by the autoluminescent *M. tuberculosis* carrying the *luxCDABE* gene cluster has been utilized as a surrogate of CFUs or visible bacterial growth for high-throughput evaluation of antimycobacterial compounds *in vitro* and in *vivo* (Li et al., 2023, Yang et al., 2015, Zhang et al., 2012). The combination of improved inhalation delivery method and autoluminescent *M. tuberculosis* may offers a promising approach for assessing drug activities.

Here, we have developed an autoluminescence-based, inhalable, time-efficient, non-invasive, and cost-effective antimicrobial assessing model that provides a valuable tool for evaluating the *in vivo* efficacy of lead compounds, frontline anti-TB medications, and therapeutic regimens, thereby offering insights crucial for clinical practice.

## MATERIALS AND METHODS

### Mycobacterial strains and culture conditions

The genetically engineered autoluminescent *M. tuberculosis* H37Rv (AlRv) strain, which harbors the *luxCDABE* gene cluster within its genome, was preserved at -80° (Yang et al., 2015). AlRv displays a consistent emission of blue-green light independently, a characteristic attributed to the enzymatic activity encoded by the *luxAB* genes. *luxCDE* genes are responsible for the continuous production of the substrate aldehyde necessary for bioluminescence. Remarkably, AlRv demonstrates drug susceptibility and growth rate comparable to that of the wild-type *M. tuberculosis* strain. The traditional quantification method of CFUs was replaced by the assessment of relative light units (RLUs), maintaining an approximate ratio of 10:1.

The AlRv was subcultured in Middlebrook 7H9 broth (Difco, Detroit, MI, USA), supplemented with 10% oleic-acid-albumin-dextrose-catalase enrichment medium (BBL, Sparks, MD, USA), 0.2% glycerol, and 0.05% Tween80 at 37°. The AlRv culture was grown until the optical density, measured at 600 nm, reached 0.6-0.8, with a corresponding RLUs level of 5 × 10^6^/mL. The optical density was measured using a V-1000 spectrophotometer (AOE, Shanghai, China), and the RLUs were quantified using the GLOMAX 20/20 luminometer (Promega, Madison, USA).

### Antimicrobials

All pharmaceutical drugs, including rifampicin (RIF), isoniazid (INH), and ethambutol (EMB) were purchased from Meilun (Dalian, China) with a minimum purity threshold of ≥ 95%. RIF and INH were solubilized in sterilized water, serving as the solvent, to attain stock concentrations of 2 mg/mL, which were subsequently diluted to 0.5 and 0.25 mg/mL, respectively. EMB was similarly dissolved into sterilized water at its original concentration of 10 mg/mL, then further diluted to 2.5 and 0.625 mg/mL. The prepared solutions of the drugs were refrigerated at 4 [until administration.

### Aerosol infection

The female BALB/c mice (Gempharmatech, Foshan, China), aged six to eight weeks, underwent an acclimatization period of five to seven days before the initiation of experimental procedures. Subsequently, all mice were exposed to 10 mL broth culture of AlRv using an inhalation exposure system (Glas-Col, Terre Haute, IN) (Yu et al., 2020, Yu et al., 2021).

### Determination of the initial bacterial number

After infection, the mice were randomly divided into treatment and control groups, each containing six mice. On the initial day of treatment administration, anesthesia was administered individually to six mice by placing them in a container with 0.5 mL of isoflurane. It took around ten seconds for the loss of consciousness. Subsequently, the live RLUs of mice were measured by aligning their chests with the detection position of the GLOMAX 20/20 luminometer. After detection of the live RLUs, six mice were euthanized via cervical dislocation and positioned on a foam board with large head pins to secure them, ensuring their abdomens were oriented upwards. The mice were sterilized by spraying 75% ethanol and dissected to obtain the lungs. The organs were then placed into 2 mL sterilized phosphate-buffered saline, washed once with phosphate-buffered saline, and homogenized using glass tissue grinders. The lung suspensions were diluted using a 10-fold serial dilution method. Subsequently, the RLUs of 0.5 mL of undiluted and 10-fold diluted lung suspensions were measured using the GLOMAX 20/20 luminometer. Based on the detected RLUs and the ratio of CFUs to RLUs, the CFUs of the lung suspension were briefly speculated upon. The 0.5 mL properly diluted lung suspension was plated on 7H10 plates containing trimethoprim, actidione, carbenicillin, and polymyxin B to prevent contamination. All plates were then incubated in a 37° thermostatic incubator for four weeks, following which the CFUs were determined.

### Chemotherapy

Each cohort of six mice underwent daily administration of either 4 mL of solvent or drug solutions utilizing the Tow Systems Nose-Only Exposure Units (Tow-Int, Shanghai, China), as previously described (Fan et al., 2022, Yang et al., 2021, Zhao et al., 2023). The device of administration is illustrated in Figure S1. Mice were subsequently secured in a holder and connected to the exposure cabinet. The drug or solvent solution was injected into the atomizer, which uses high-frequency oscillation technology through microporous screens to generate nebulized aerosols with diameters ranging from 1 to 5 micrometers for inhalation. The drug exposure duration for each group was approximately 20-25 minutes until 4 mL of the solutions had been administered.

The study consists of two main experiments, delineated as Experiment 1 and Experiment 2. The experimental protocols remained largely consistent, with the primary variations involving alterations in pharmacological dosages and the timing of medication administration subsequent to infection. Detailed protocols for Experiments 1 and 2 were provided in Tables 1 and 2, respectively. The live RLUs of the solvent group were continuously monitored from the day 11 for Experiment 1 (or day 13 for Experiment 2) post-infection. Measurements of RLUs for all groups were performed once the live RLUs of the solvent-treated mice exceeded 400. Based on our experiences, live RLUs exceeding 400 indicated a higher level of lung RLUs and CFUs, suitable for evaluating drug efficacy. Treatment of mice was discontinued upon reaching live RLUs exceeding 400. The total duration of administration for Experiments 1 and 2 was 14 and 15, respectively. Subsequently, all mice were euthanized the following day to determine lung RLUs and CFUs following the method mentioned above. The plates were then incubated at 37° for four weeks, and then CFUs were determined.

**Table 1.** Scheme of experiment 1.

| Drug/Dosage<sup>1</sup><br>( mg/mL ) | Number of mice sacrificed at the given time points |  |  |  | Total |
| --- | --- | --- | --- | --- | --- |
|  | D0<sup>2</sup> | D3<sup>3</sup> | D11-17 | D18<sup>4</sup> |  |
| Solvent |  | 6 | Detecting chest | 6 | 12 |
| RIF 2 |  | Treatments | RLUs of live mice | 6 | 6 |
| INH 2 |  | initiation | in solvent group | 6 | 6 |
| EMB 10 | Infection |  | daily from day 11 | 6 | 6 |
|  |  |  | until live RLUs |  |  |
|  |  |  | exceed 400 |  |  |
| Total | 30 | 6 |  | 24 | 30 |

<sup>1</sup> RIF, rifampicin; INH, isoniazid; EMB, ethambutol; Solvent, sterilized water; the number after the drug represents the dosage.
<sup>2</sup> D0, a total of 30 mice were infected with AIrV by aerosol.
<sup>3</sup> D3, the mice were randomly divided into groups and treated daily from the day 3 to day 16. The live RLUs, the lung suspension RLUs and CFUs of six mice were detected before the treatment initiation.
<sup>4</sup> D18, the mice were sacrificed and the lungs were removed for detecting RLUs and CFUs.
All CFUs were counted 4 weeks after incubation at 37°C.

**Table 2.** Scheme of experiment 2.

| Drug/Dosage<sup>1</sup><br>( mg/mL ) | Number of mice sacrificed at the given time points |  |  |  | Total |
| --- | --- | --- | --- | --- | --- |
|  | D0<sup>2</sup> | D1<sup>3</sup> | D13-16 | D17<sup>4</sup> |  |
| Solvent |  | 6 |  | 6 | 12 |
| RIF 2 |  |  |  | 6 | 6 |
| RIF 0.5 |  |  | Detecting chest | 6 | 6 |
| RIF 0.125 |  |  | RLUs of live mice | 6 | 6 |
| INH 2 | Infection | Treatments<br>initiation | in solvent group | 6 | 6 |
| INH 0.5 |  |  | daily from day 13 | 6 | 6 |
| INH 0.125 |  |  | until live RLUs | 6 | 6 |
| EMB 10 |  |  | exceed 400 | 6 | 6 |
| EMB 2.5 |  |  |  | 6 | 6 |
| EMB 0.625 |  |  |  | 6 | 6 |
| Total | 66 | 6 |  | 60 | 66 |

<sup>1</sup> RIF, rifampicin; INH, isoniazid; EMB, ethambutol; Solvent, sterilized water; the number after the drug represents the dosage.
<sup>2</sup> D0, a total of 66 mice were infected with AIRv by aerosol.
<sup>3</sup> D1, the mice were randomly divided into groups and treated daily from day 1 to day 15. The live RLUs, the lung suspension RLUs and CFUs of six mice were detected before the treatment initiation.
<sup>4</sup> D17, the mice were sacrificed, and the lungs were removed for detecting RLUs and CFUs.
All CFUs were counted 4 weeks after incubation at 37°C.

### Statistical analysis

Log-transformed data using GraphPad Prism version 8.3.0 was used to determine drug efficacy through a two-way analysis of variance, with *P* < 0.05 as statistical significance.

## RESULTS

### The results of experiment 1

In experiment 1, the initial CFUs of the infected bacteria were approximately 4000. The live RLUs of all treatment groups were significantly lower than those of the solvent group, with all differences exceeding 200 (Figure 1, *P* < 0.0001). Additionally, these live RLUs were comparable to those of the before-treatment (BT) group (Figure 1, *P* > 0.05). Furthermore, the lung RLUs of all drug-treated groups were substantially lower than those of the solvent group, with all log_10_RLUs/lung differences exceeding 1.5 (Figure 1, *P* < 0. 0001). Specifically, the lung RLUs of the RIF or EMB-treated groups were nearly identical to those of the BT group (Figure 1, *P* > 0.05), while the INH-treated group exhibited slightly lower lung RLUs than the BT group (Figure 1, *P* < 0.05). Moreover, the lung CFUs of all treatment groups were markedly lower than that of the solvent group, as all differences were over 1.5 (Figure 1, *P* < 0.0001). The RIF group had higher lung CFUs, whereas the INH group had lower lung CFUs compared to the BT group (Figure 1, *P* < 0.01 and *P* < 0.0001, respectively). The lung CFUs of the EMB-treated group were almost similar to those of the BT group (Figure 1, *P* > 0.05). The data from experiment 1 indicated that the autoluminescence-based inhalation delivery method can be utilized to establish a non-invasive model for testing drugs activities *in vivo*, using the same batch of live mice within 18 days after infection. Furthermore, the effectiveness of drugs can be further evaluated as either bacteriostatic or bactericidal by comparing lung CFUs of the different treatment groups rather than RLUs, as lung RLUs of the BT group were close to the background values.

**Figure 1.**
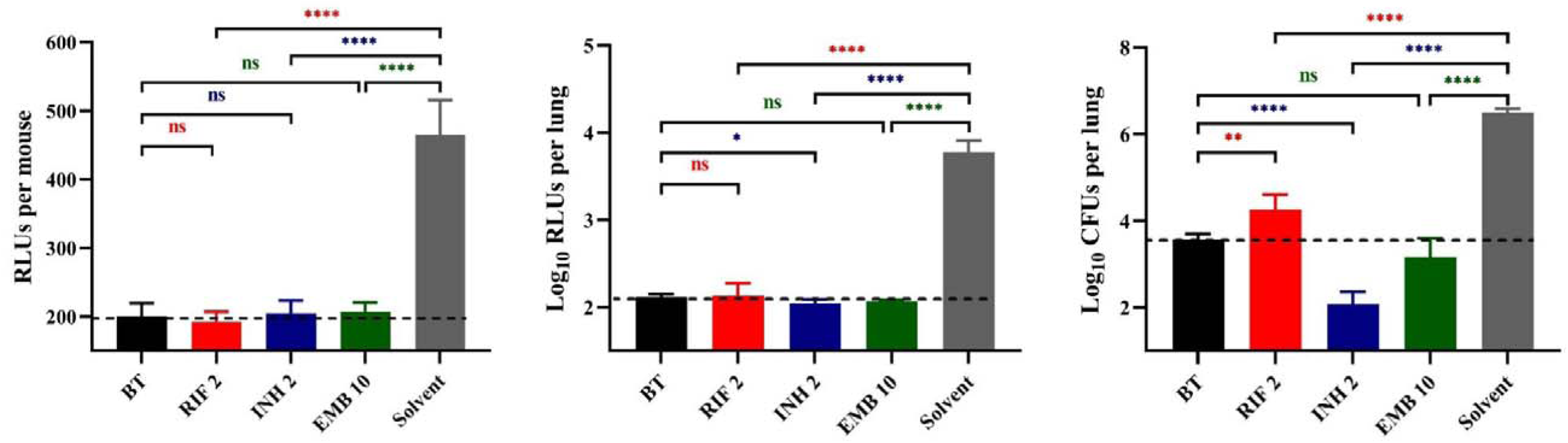
Anti-TB drug activities by inhalation administration in Experiment 1. BT, before treatment; RIF, rifampicin; INH, isoniazid; EMB, ethambutol; Solvent, sterilized water; the number after the drug represents the dosage; *, *P* < 0.05; **, *P* < 0.01; ***, *P* < 0.001; ****, *P* < 0.0001; ns, *P* > 0.05, no significance. The dotted lines represent the background values.

### The results of experiment 2

Based on the results of Experiment 1, two additional dosages of each drug were added in Experiment 2. The initial CFUs of the infected bacteria in Experiment 2 were found to be roughly equivalent to those recorded in Experiment 1. The live RLUs of the groups treated with RIF 2, INH 2, INH 0.5, EMB 10, EMB 2.5, and EMB 0.625 mg/mL were significantly lower compared to the solvent group (Figure 2a, *P* < 0.0001, *P* < 0.0001, *P* < 0.01, *P* < 0.001, *P* < 0.001, and *P* < 0.05, respectively). Conversely, the live RLUs of RIF 0.5, RIF 0.125 and INH 0.125 mg/mL groups did not show significant differences from those of the solvent group (Figure 2a, *P >* 0.05). These findings were also largely supported by both the lung RLUs and CFUs (Figure 2b and 2c), except for the lung CFUs of the RIF at 0.5 mg/mL group, which were slightly significantly lower than that of the solvent group (Figure 2c, *P* < 0.05). In summary, these findings suggest a negative correlation between live RLUs and administered dosages, indicating that live RLUs could serve as an initial criterion for assessing the therapeutic efficacy.

**Figure 2.**
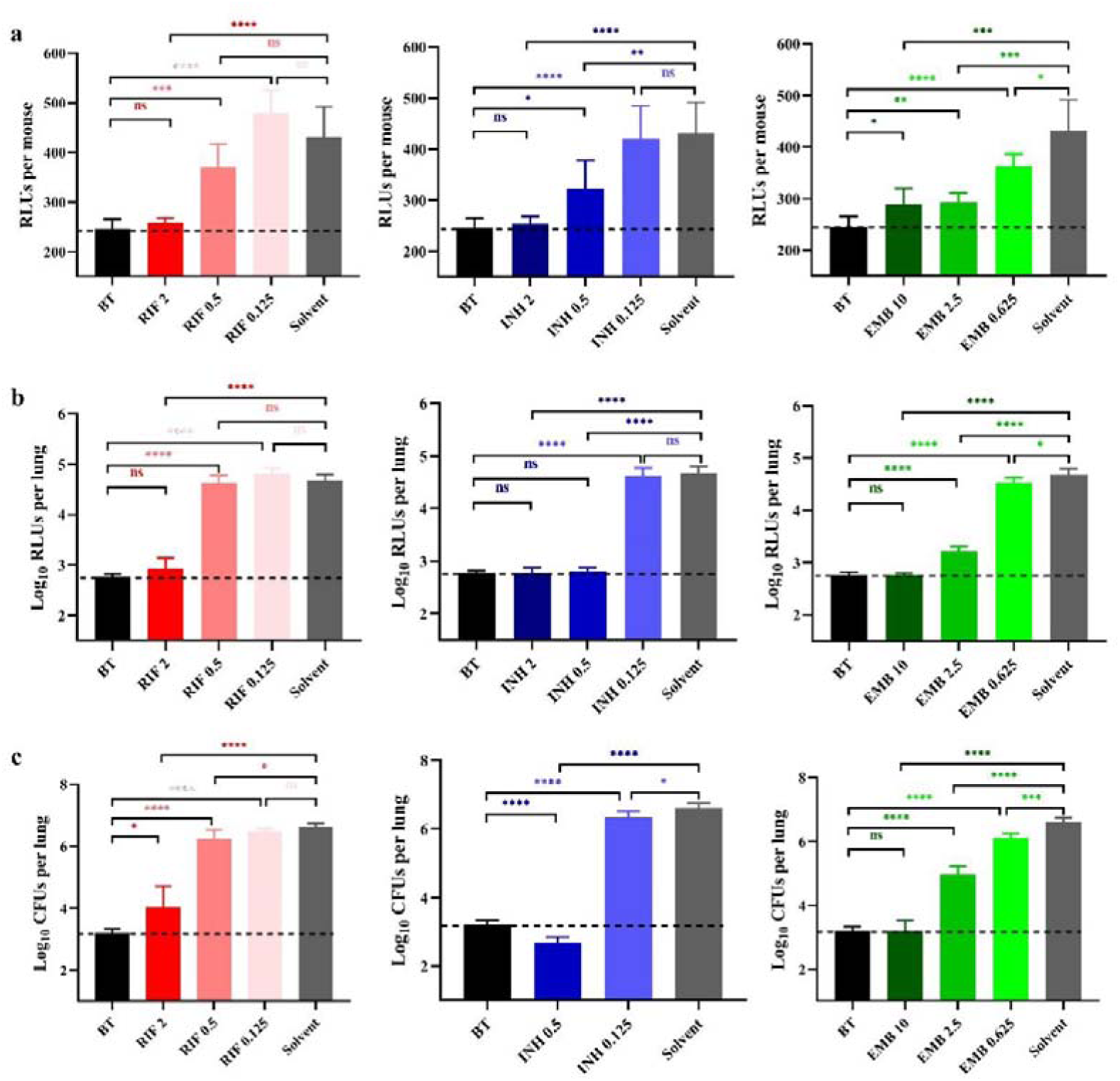
Anti-TB drug activities by inhalation administration in Experiment 2. (a) Light intensities of live mice. (b) The suspension light intensities. (c) The lung *M. tuberculosis* burden. BT, before treatment; RIF, rifampicin; INH, isoniazid; EMB, ethambutol; Solvent, sterilized water; the number after the drug represents the dosage; *, *P* < 0.05; **, *P* < 0.01; ***, *P* < 0.001; ****, *P* < 0.0001; ns, *P* > 0.05, no significance. The dotted lines represent the background values.

The lung RLUs of the RIF 2 mg/mL group demonstrated a significant reduction compared to those in the solvent group, with a log_10_RLUs/lung difference exceeding 1.5 and even similar to that of the BT group (Figure 2b, *P* < 0.0001 and *P* > 0.05, respectively). Conversely, the lung RLUs in the RIF 0.5 and 0.125 mg/mL groups were comparable to that in the solvent group, indicating a lack of evident anti-*M. tuberculosis* activity at these dosages (Figure 2b, *P* > 0.05). The lung RLUs of the INH 2 and 0.5 mg/mL groups were significantly lower than that of the solvent group (Figure 2b, *P* < 0.0001), as all differences in log_10_RLUs/lung exceeded 1.5 and were comparable to that of the BT group (Figure 2b, *P* > 0.05). The lung RLUs of the INH 0.125 mg/mL group were similar to those of the solvent group (Figure 2b, *P* > 0.05).

In comparison, the lung RLUs of the EMB 10 and 2.5 mg/mL groups exhibited a pronounced decrease compared to those in the solvent group, with log_10_RLUs/lung differences exceeding 1.5 and 1, respectively (Figure 2b, *P* < 0.0001). Meanwhile, the lung RLUs of the EMB 10 mg/mL group were equivalent to that of the BT group (Figure 2b, *P* > 0.05), whereas the lung RLUs in the EMB 0.625 mg/mL group showed less significant differences from the solvent group (Figure 2b, *P* < 0.05). In conclusion, the lung RLUs results are consistent with those obtained from the live RLUs, providing further support for the utility of detecting live RLUs in determining drug activity.

The lung CFUs in the RIF 2 mg/mL group exhibited a significant reduction compared to those in the solvent group with a difference in log_10_CFUs/lung exceeding 1.5 (Figure 2c, *P* < 0.0001). The lung CFUs in the RIF 0.5 and 0.25 mg/mL groups were approximately comparable to the solvent group (Figure 2c, *P* < and close to 0.05 and *P* > 0.05, respectively). Notably, no colonies were observed on the plates of the INH 2 mg/mL group, indicating possibly eradication of AlRv from the mice lungs, considering that only a fraction of the lung suspension from each mouse was plated. The CFUs of the INH 0.5 mg/mL group were also significantly lower than that of the solvent group, with a difference of log_10_CFUs/lung exceeding 1.5 and even lower than the BT group (Figure 2c, *P* < 0.0001). Moreover, all EMB dosages resulted in significantly reduced lung CFUs compared to the solvent control (Figure 2c, *P* < 0.0001, *P* < 0.0001, and *P* < 0.001, respectively). However, only the log_10_CFUs/lung difference between the EMB 10 mg/mL and solvent group exceeded 1.5, with the lung CFUs in the EMB 10 group being nearly identical to those in the BT group (Figure 2c, *P* > 0.05). These CFU results were consistent with the findings derived from live or lung RLUs data. Overall, based on the comprehensive experimental data presented, we have successfully established an autoluminescence-based inhalation administration murine model for evaluating drugs activities against *M. tuberculosis in vivo*.

## DISCUSSION

*M. tuberculosis* imposes a substantial burden, particularly in developing countries (Chong and Lim, 2009, Grüber, 2020). Despite ongoing efforts, the development of effective anti-*M. tuberculosis* drugs remains confronted with numerous challenges, including the complexity of evaluation process and the emergence of drug-resistant strains (McGrath et al., 2014, Olaru et al., 2015). Over the past several decades, inhalable therapy has emerged as a promising approach for treating pulmonary infections, offering improved cure rates and bacterial eradication (Riveiro et al., 2023). However, the evaluation of inhalable drugs still presents challenges such as complex manipulation and time-consuming results completion. Recent researches have revealed that a recombinant autoluminescent strain of *M. tuberculosis* carrying *luxCDABE* gene cluster facilitates rapid, non-invasive, real-time monitoring of the RLUs in live mice during experimental chemotherapy, providing a valuable tool for assessing treatment efficacy (Yang et al., 2015, Zhang et al., 2012). Hence, we have developed a novel murine model by combining the improved inhalation method and autoluminescent *M. tuberculosis*.

As of the present, four primary research reports, encompassing three inhalation administration models, have explored the detection of anti-TB drug efficacy through airway administration (Garcia-Contreras et al., 2007, Garcia-Contreras et al., 2010, Gonzalez-Juarrero et al., 2012, Verma et al., 2013). Among these studies, only two have employed murine models for their investigations. Table S1 presents a comprehensive comparison of the advantages and disadvantages of each report relative to our study. Compared to these existing inhalation administration model, our innovative method achieves several improvements. Firstly, it is cost-effective and user-friendly, employing mice instead of larger animals. Secondly, it eliminates the need for anesthesia in mice during administration, simplifying the instrument operation. We also attempted to employ published administration method in mice infected with AlRa via tail vein injection, but encountered significant challenges in administering drugs daily due to the mice developing swollen throats (Gonzalez-Juarrero et al., 2012). However, we did observe that mice treated with INH exhibited reduced RLUs compared to both the untreated control and the day before administration (unpublished data). Thirdly, the results are relatively objective and repeatable, minimizing human factors. Fourthly, the same treatment group, consisting of six mice, is administrated daily and simultaneously. In fact, we indeed succeeded in administrating the same six mice 2-3 times/day and even with different drugs. Additionally, our method is non-invasive, measuring the live RLUs of mice as a surrogate marker for CFUs for drug *in vivo* activity testing. Lastly, it reduces the time required for obtaining results from several months to 16 or 17 days.

We have evaluated the *in vivo* efficacy of three first-line TB drugs using our model, including RIF, INH, and EMB. Our model can accelerate assessment of drug activity, with results obtainable within 17 or 18 days or even shorter by detecting either live RLUs of mice or RLUs of lung suspensions. Our findings demonstrate that RIF, INH, and EMB, at the dosage of 0.5, 0.5, and 0.625 mg/mL, exhibited significant anti-*M. tuberculosis* activity *in vivo*. Notably, INH exhibits potential for complete eradication of lung bacteria at increased dosage up to 2 mg/mL in the current model. Moreover, the CFUs results corroborate the findings inferred from live RLUs, affirming the utility of live RLUs as a surrogate marker for CFUs in mice. This innovative model holds promise for enhancing the assessment of anti-*M. tuberculosis* drug efficacy *in vivo*, thereby reducing the likelihood of overlooking potentially effective treatments and minimizing the risk of adverse effects.

The drug’s activity can be determined by comparing the lung suspension RLUs of different groups within a shorter duration than the current 17-day period (from infection to get the results), since the lung RLUs become significantly higher than the background value a few days earlier than the live RLUs do. Furthermore, drug efficacy could potentially be observed within a considerably shorter time frame in live mice if mice infected with high dose AlRv via tail vein injection route, similar to the previous studies showed that the live RLUs could be detected at the day after infection, although this may result in a higher standard deviation (Liu et al., 2019, Olaru et al., 2015, Zhang et al., 2012). Further investigation is warranted to determine whether other anti-TB drugs or potent compounds demonstrate efficacy using our model. Additionally, exploring alternative formulations or solvents may further enhance the drug’s effectiveness of drugs beyond sterilized water.

In conclusion, our study establishes an autoluminescence-based, inhaled, and non-invasive administration murine model for evaluating anti-*M. tuberculosis* drug *in vivo* efficacy. This model holds promise for expediting the development of innovative anti-*M. tuberculosis* drugs and treatment regimens, addressing the urgent need for more effective TB therapies. Meanwhile, this improved inhalation model can also be used for developing drugs targeting other diseases and assessing the toxicity of compounds.

## DATA AVAILABILITY STATEMENT

The original contributions presented in the study are included in the article, further inquiries can be directed to the corresponding authors.

## ETHICS STATEMENT

The animal care and experimental protocols were approved by the Laboratory Animal Ethics Committee of the Guangzhou Institutes of Biomedicine and Health, Chinese Academy of Sciences.

## AUTHOR CONTRIBUTIONS

Conceived the project: Tianyu Zhang, Jinxing Hu, Xirong Tian, Yamin Gao; Designed the research: Tianyu Zhang, Jinxing Hu, Xirong Tian, Yamin Gao; Performed the studies: Xirong Tian, Yamin Gao, Jingran Zhang, Yanan Ju, Jie Ding, Sanshan Zeng, Wanli Ma; Interpreted the results: Tianyu Zhang, Xirong Tian, Yamin Gao; Drafted the manuscript: Tianyu Zhang, Jinxing Hu, Xirong Tian, Yamin Gao, H.M. Adnan Hameed, Nanshan Zhong, Gregory M. Cook; Final approval of the manuscript: all authors.

## FUNDING

This work was supported by the National Key R&D Program of China (2021YFA1300900; 2023YFFO713605), the National Natural Science Foundation of China (81973372, 21920102003, 82061138019, 32300152), Guangdong Provincial Basic and Applied Basic Research Fund (2022A1515110505), Guangzhou Science and Technology (2024A04J4273), Chinese Academy of Sciences (CAS)-President’s International Fellowship Initiative (2023VBC0015), China Postdoctoral Science Foundation (2022M723164), the Chinese Academy of Sciences Grants (YJKYYQ20210026), State Key Laboratory of Respiratory Disease (SKLRD-Z-202301, SKLRD-Z-202414), and Guangzhou Science and Technology Planning Project (2023A03J0992).

## Notes

### Competing Interest Statement

The authors have declared no competing interest.

### Summary of Updates

Supplemental files added; Abstract, Introduction and Discussion modified.

