## Supplementary material for "Establishment of an Inhalation Administration Non-invasive Murine Model for Rapidly Testing Drug Activity against *Mycobacterium tuberculosis*": Figure S1 and Table S1

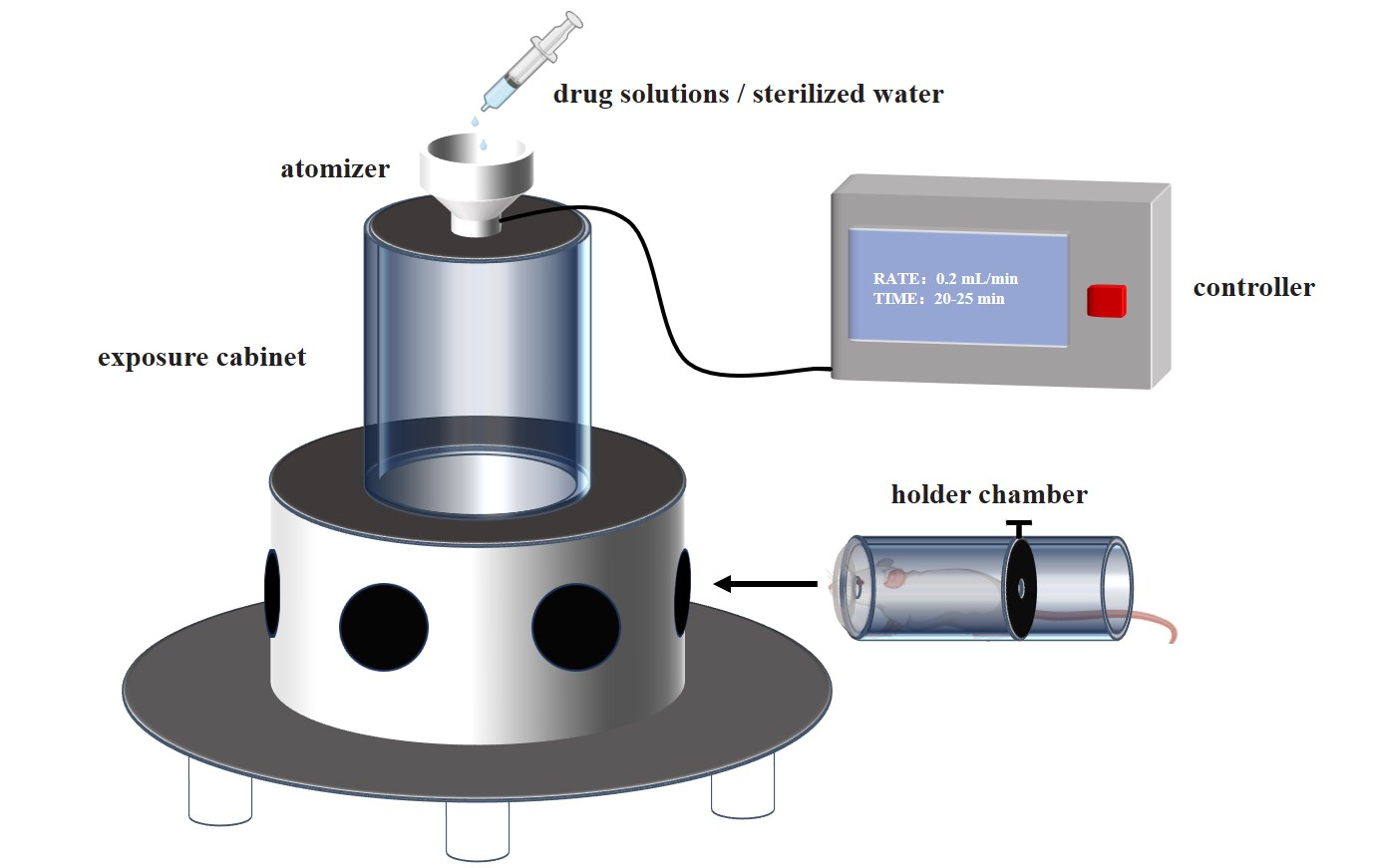

**Figure S1. Schematic diagram of inhalation administration using the Tow Systems Nose-Only Exposure Units.**

**Table S1. Summary of comparative strengths and weaknesses of the inhaled administration studies.**

| Animal | Anesthesia | Standardization  /Repeatability | No. of animals administered each time | Operability | Drug amount | Administration speed | Administration frequency | Live detection of activities | Time to get results after administration of the last dose |
| --- | --- | --- | --- | --- | --- | --- | --- | --- | --- |
| Mouse**^(1)^**^*^ | Yes | Low | 1 | Hard | Small | ~15 min | 3/week | No | 3-5 weeks |
| Guinea pig**^(2, 3)^** | - | High | 4 | Hard | Media | 30 min/low-dose or 60 min/ high-dose | daily | No | 3-5 weeks |
| Mouse**^(4)^** | - | High | 1 | Hard | Small | - | 2/week | No | 3-5 weeks |
| Mouse**^(this study)^** | No | High | 6 | Easy | Large | 20~25 min | One to several times/day | Yes | 1 day |

^*^ We also tried this method and found the throat of the mouse showed somewhat swelling after administration, so it was hard or even almost impossible to be administer the same mouse daily.

-, unclear.
